# An anatomical substrate of credit assignment in reinforcement learning

**DOI:** 10.1101/2020.02.18.954354

**Authors:** J Kornfeld, Y Wang, M Januszewski, A Rother, P Schubert, M Goldman, V Jain, W Denk, MS Fee

## Abstract

A key problem in learning is credit assignment. Biological systems lack a plausible mechanism to implement the backpropagation approach, a method that underlies much of the dramatic progress in artificial intelligence. Here, we use automated connectomic analysis to show that the synaptic architecture of songbird basal ganglia (Area X) supports local credit assignment using a variant of a node perturbation algorithm proposed in a model of reinforcement learning. Using two volume electron microscopy (vEM) datasets, we find that key predictions of the model hold true: axons that encode exploratory variability terminate predominantly on dendritic shafts, while axons that encode song timing (context) terminate predominantly on spines. Based on the detailed EM data, we then built a biophysical model of reinforcement learning that suggests that the synaptic dichotomy between variability and context encoding axons facilitates efficient learning. In combination, these findings provide strong evidence for a general, biologically plausible credit assignment model in vertebrate basal ganglia learning.

**One Sentence Summary:** Using automated connectomic analysis and biophysical modeling, we show how the basal ganglia could solve the credit assignment problem on the synaptic level.

## Main text

Neural circuits that control decisions and actions are recurrently connected and involve many network layers from sensory inputs to motor output. Yet, as we learn, some mechanism specifies precisely which synapses, out of trillions, are to be modified and in what way. The backpropagation algorithm ^1^ is powerful because it directly calculates, based on the network architecture, the derivative of output errors with respect to every synaptic weight, providing an efficient method to update synaptic strengths. However, it remains unclear whether backpropagation or its variants are biologically implemented, or even plausible ^2–4^. An alternative approach to implement gradient-based learning is weight- or node-perturbation ^5,6^, in which the activity of a specific synapse or neuron is stochastically varied to determine its contribution to the output. Here we use a connectomic approach to study the biological implementation of stochastic gradient descent, which requires as-yet unknown circuit structures to inject variability, correlate variability with reward signals, and correctly assign credit to relevant synapses.

In this sense, node perturbation is conceptually similar to behavioral trial-and-error reinforcement learning (RL) ^7,8^. In the vertebrate brain, RL involves the basal ganglia ^9^, where a multitude of sensory and other context (or state) signals converge with action and outcome signals to determine which actions in each state lead to the best outcomes. Songbird Area X, the basal ganglia circuit dedicated to song learning ^10^, receives synaptic input from two key cortical areas ^11^: LMAN (lateral magnocellular nucleus of the anterior neostriatum), which acts as a source or vocal variability in the song motor system ^12,13^, and HVC (proper noun), which generates a sparse timing signal that controls vocal output at each moment in the song ^14^. Area X also receives information about song performance via dopaminergic (DA) axons originating in the ventral tegmental area (VTA) ^15^ (Fig. 1A). Fee and Goldberg proposed ^16^ that medium spiny neurons (MSNs) in Area X integrate signals from HVC, LMAN, and VTA to detect which song variations (from LMAN) at which times in the song (from HVC) lead to improved song performance (from VTA). The model posits that convergence of these factors drives plastic changes in HVC-MSN synapses within the basal ganglia such that HVC drives improves song performance at each moment in subsequent song renditions ^16^.

**Fig. 1.**
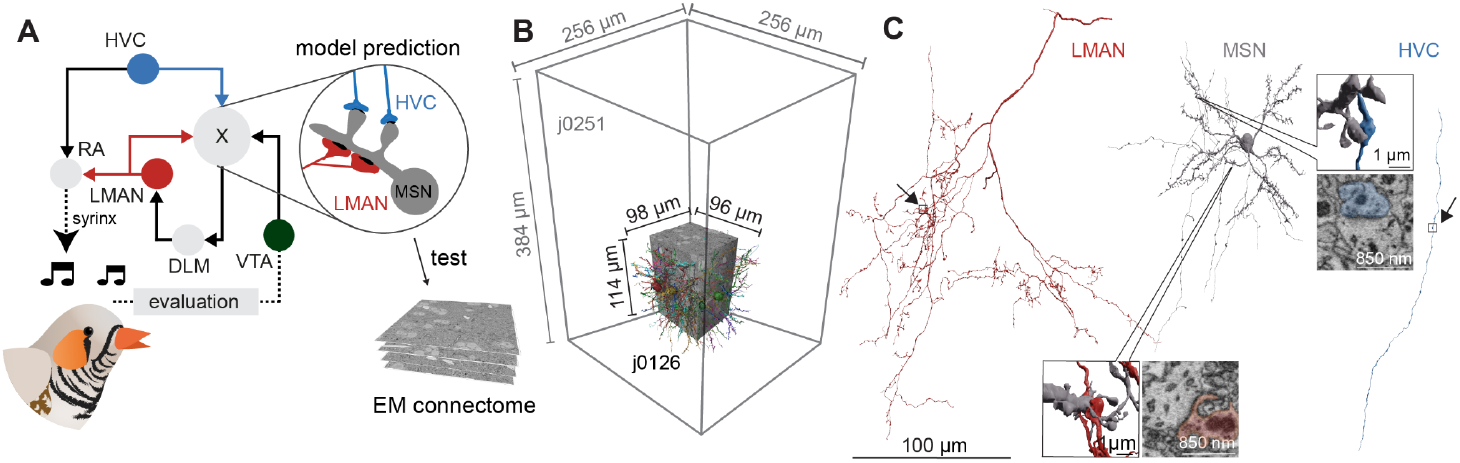
Anatomy of the zebra finch song system and Area X connectome analysis. (A) Key brain regions and their connections involved in zebra finch song learning and generation; solid lines indicate mono-synaptic connections; broken lines, multi-synaptic. (B) Two vEM data sets of adult male zebra finches were analyzed for this study: j0126 and j0251. (C) Typical neurites from putative corticostriatal axons HVC, LMAN, and a MSN from j0251 (3D visualization). Scale bar for HVC, LMAN and MSN 100 µm. LMAN-MSN synapse data set coordinates: 11370, 7776, 8999, HVC - MSN synapse: 9920, 5976, 10514.

At a synaptic level, precise temporal coincidence of HVC and LMAN input onto MSN dendrites is hypothesized to form a transient biochemical eligibility trace ^17^ at the HVC-MSN synapse. This tags the synapse as a candidate for strengthening, conditional on a subsequent DA signal, indicating improved song performance ^6,18–20^, similar to three-factor learning rules ^21,22^. Spiny synapses are well suited to carry out the proposed functions of the HVC-MSN connection: they are electrically and biochemically compartmentalized ^23^, can function as coincidence detectors ^24,25^, exhibit robust synaptic plasticity ^26^, and have been proposed as loci of eligibility traces ^17^. In contrast to the HVC inputs, the model posits that LMAN synapses are neither plastic, nor carry an eligibility trace, but rather signal the occurrence of actions globally across MSN dendrites—a function possibly more suitable to shaft synapses. Altogether, these observations lead to a specific prediction (Fig. 1A) that HVC-MSN synapses preferentially terminate on spines, while LMAN-MSN synapses preferentially terminate on dendritic shafts ^19^.

To examine the anatomical fine structure of inputs to MSN dendrites and test these predictions, we acquired two data sets by serial block-face electron microscopy (SBEM) ^27^ from Area X of two adult (>120 dph) male zebra finches. The neural circuitry contained in the EM volume was then reconstructed using largely automated methods for neurite segmentation, synapse identification and morphology classification ^28–31^ (Fig. 1 B, C). The machine output of the smaller data set (j0126), was inspected and improved by manual proof-reading (see Methods, <1000 hours). However, later inspection revealed that our conclusions are the same for analyses carried out on fully automated reconstructions, without manual neurite proofreading (see Supplementary Text and Fig. S2). Based on visual inspection, we excluded only obviously erroneous reconstructions for the larger data set (j0251, see companion paper by Rother et al.).

To interpret the synaptic architecture associated with HVC and LMAN axons onto MSN dendrites, we needed to identify these cells in the dataset. We used a neural network-based morphology classifier ^30,31^ trained on the basis of known morphological characteristics. These characteristics included the rate of branching, which is especially low for HVC axons, the presence of a dense terminal plexus, typical of LMAN axons ^32,33^, and the density of spines on dendrites, to identify MSNs ^34^ (Fig. 1C, Fig. S1).

We started by analysing the nature of HVC and LMAN synaptic contacts on MSN dendrites in the j0251 data set (Fig. 2A). Together these inputs contribute >90% of all excitatory synapses onto MSNs. On average MSNs receive 315+-100.8 (sd) HVC synapses and 34+-13.8 (sd) LMAN synapses. HVC synapses contribute eightfold more synaptic area than LMAN synapses (Fig. 2B), a ratio much larger than the 1-to-1 ratio between the numbers of axons (about 10,000) entering from each region ^35,36^, but smaller than the 10-to-1 ratio between the total axon path lengths from these inputs regions (∼49 m vs. ∼5 m, in the j0251 data set).

**Fig. 2.**
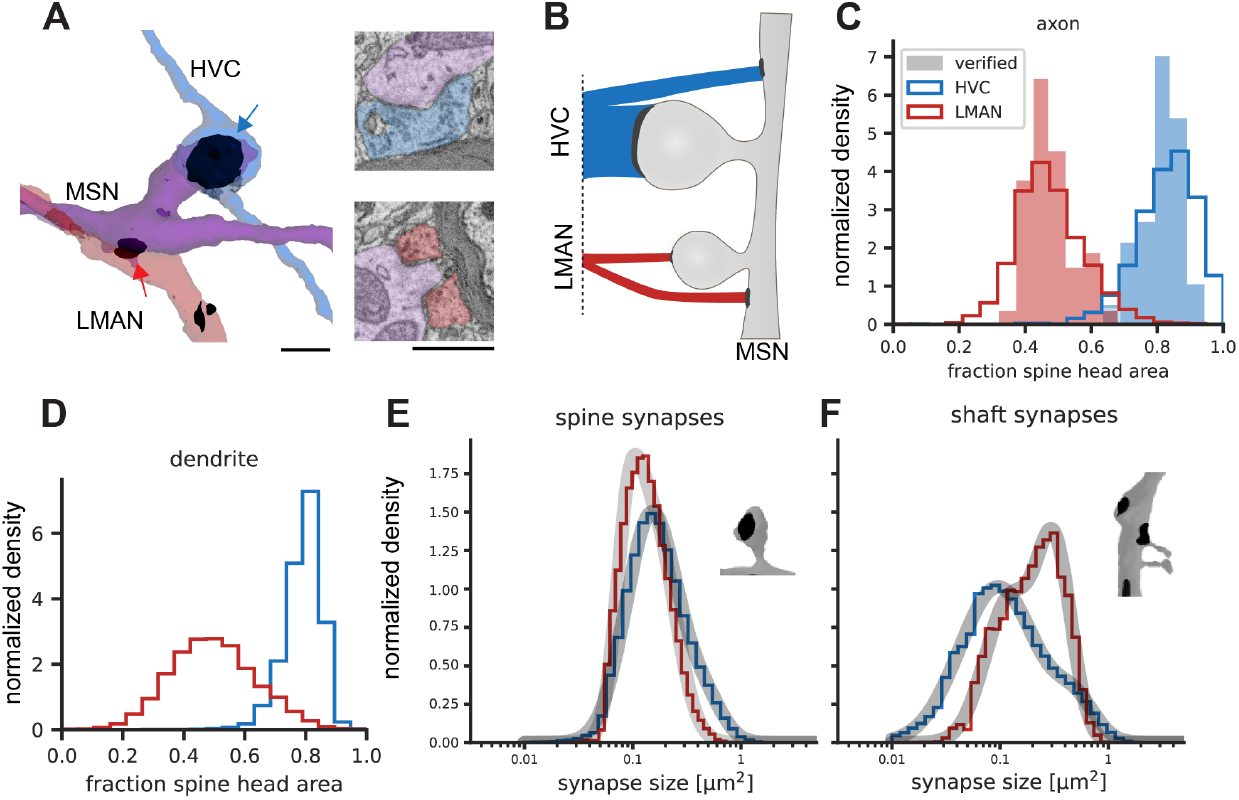
Spine / shaft preference of HVC and LMAN inputs to MSN dendrites. Surface rendering (A, left) and cross section (right), both showing the same micro volume from the j0126 data set containing one HVC-MSN (blue arrow) and one LMAN synapse (red arrow), which contact, respectively, a spine head and the dendritic shaft of the same MSN (purple). Scale bars 1 µm. (B) Conceptual sketch of the ratio between spine- and shaft bound-fractions for HVC (top) and LMAN (bottom) inputs and the contribution of each of the four cases to the total excitatory synaptic area on MSN dendrites. (C) Normalized histogram of fraction of spine head area per neurite for different axon types with MSN dendrites. Verified axons were manually inspected and used as classifier ground truth. (D) Analysis from the perspective of MSNs, i.e. each data point underlying the two distributions is the fraction of synaptic input area on spines one MSN receives, from LMAN or HVC synapses, respectively. (E, F) Distribution of synapse size for spine and shaft synapses of HVC and LMAN synapses with MSNs. Also shown are fits (grey curves) to a gaussian mixture model with two truncated log-normal distributions (Methods). Data in C-F from j0251, see Fig. S2 for j0126 data.

Consistent with the model’s prediction, HVC axons synapse preferentially with MSN spines, while LMAN axons synapse preferentially with MSN shafts. In the larger dataset, quantitative analysis of synaptic areas showed that 78.8% of HVC input onto MSNs, but only 46.7% of LMAN input onto MSNs, was found on spines. In contrast, the synaptic areas of HVC and LMAN axons terminating on MSN dendritic shafts were 18.0% and 50.0%, respectively. (Less than 3.3% of HVC and LMAN axon synaptic area terminated on MSN compartments other than spine or dendritic shaft.) An analysis of manually classified HVC and LMAN axons yielded similar results (84.0% vs 46.5% onto spines and 16.0% vs 53.5% onto shafts; 53 HVC and 57 LMAN axons, respectively). The results from the smaller data set, obtained from a different adult male zebra finch, yield essentially identical results for HVC and LMAN axons: 86.7% vs 44.7% of the synaptic area was onto spines (see Supplementary Information for details and replication of other analyses). In comparison to the overall synaptic area distributions (spine vs shaft) of HVC and LMAN axons, these fractions correspond to a strong enrichment of LMAN inputs onto shaft synapses (by a factor of 2.4 and 3.1 for the large and small data set, respectively), and a smaller enrichment of HVC inputs onto spines (4% and 5%, respectively, see Methods). We found no evidence for subpopulations of HVC or LMAN axons with distinct spines/shaft ratios; a similar analysis also revealed no distinct subpopulations of MSN dendrites on this basis (Fig. 2C, D).

Given the very different roles for HVC and LMAN synapses postulated in the model, we wondered if these synapses also differ in their distribution of synaptic areas. We found that HVC synapses onto spines were larger in area than those from LMAN (mean = 0.21 µm^2^ vs 0.15 µm^2^, median = 0.16 µm^2^ vs 0.13 µm^2^, n = 1,920,408 and 141,494 for HVC and LMAN, respectively), while LMAN synapses onto MSN shafts were larger than those from HVC (mean = 0.24 µm^2^ vs 0.17 µm^2^, median = 0.21 µm^2^ vs 0.11 µm^2^, n = 95,757 and 521,401, for LMAN and HVC, respectively). These differences were highly significant (two-sided Mann–Whitney U tests; all p-values < 0.01). In summary, the larger size of HVC spine synapses, compared to those from LMAN, is consistent with their hypothesized role in driving temporally specific MSN activity ^37^, while the larger size of LMAN shaft synapses is consistent with their hypothesized role in depolarizing MSN dendrites to signal LMAN-driven song variations.

We noted that the size distribution of each of these synapse classes (HVC-spine, HVC-shaft, LMAN-spine, LMAN-shaft) did not appear to be simple log-normal, but rather followed the shape of a log-normal mixture (Fig. 2E,F, Fig. S2), reminiscent of a finding of bimodal synapse sizes in the mammalian cortex ^38^. Interestingly, in our data, the bimodality is more pronounced for excitatory shaft synapses than for spine synapses (Fig. 2F, Fig. S2).

To quantitatively assess whether the observed neural architecture can implement credit assignment during vocal learning, we constructed a biophysical simulation of a neural circuit model incorporating our anatomical findings. Based on the 3D reconstruction of a complete MSN (Fig. 3A, left), we built a multi-compartment biophysical model with realistic morphology and embedded it in a simple reinforcement learning circuit (Fig. 3A, right; see Methods). As previously proposed ^39^, the learning is implemented via the convergence of three inputs to the MSN: HVC spikes within a 10 ms note of the song, LMAN spikes representing collateral discharge of motor exploration, and a dopaminergic signal encoding reward prediction error (see Methods). The HVC input targets the MSN spine heads whereas the LMAN input targets the MSN dendritic shaft (Fig. 3B). The key mechanism by which the model implements credit assignment is that the LMAN input onto the dendritic shaft generates strong depolarization that spreads into nearby spine heads (Fig. 3C). When the LMAN input follows the HVC input with an appropriate delay around 15 ms ^40,41^, this depolarization relieves the magnesium block of the NMDA receptors, triggering a large calcium influx (Fig. 3D) that activates an eligibility trace for synaptic plasticity ^42–45^. If LMAN-driven variability improves song performance, this leads to the release of dopamine that conveys a positive reward prediction. The dopamine signal, together with the eligibility trace, triggers plasticity that strengthens the HVC-MSN synapses. As a result, in future renditions of the song, HVC input will increase the firing of the MSN at that time. This, through a pathway through the motor thalamic nucleus DLM (Fig. 3A, right, dashed line), drives spiking in the LMAN neuron (Fig. 3E), leading to a learned improvement of the song.

**Fig. 3.**
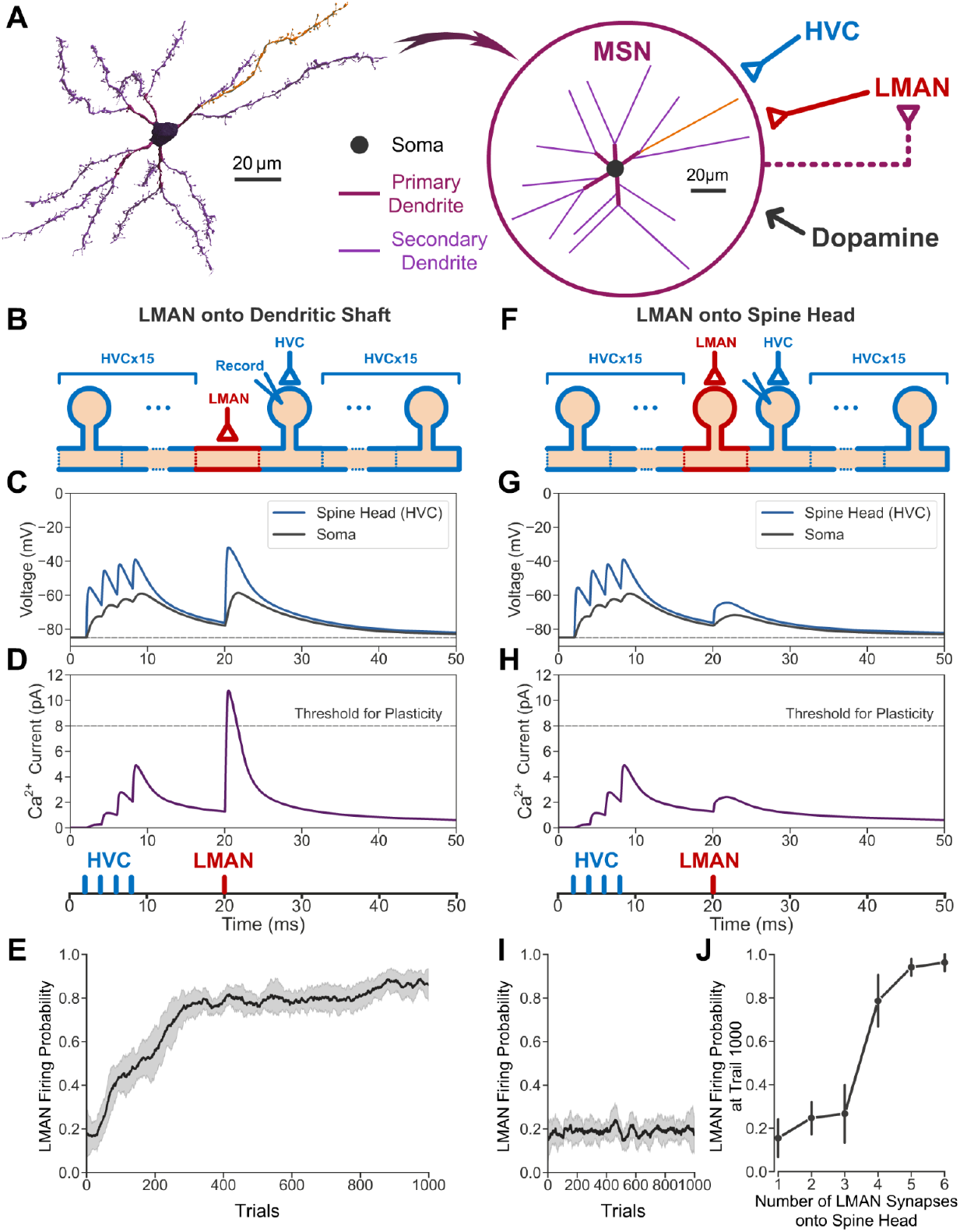
Biophysical model of credit assignment in reinforcement learning. (A) Left: 3D reconstruction of a complete MSN. The axon is hidden for clarity. Right: Morphology of the model MSN and diagram of the neural circuit model of reinforcement learning. Spines of the MSN (not shown) are equidistant on the secondary dendrites with a density consistent with our data (one spine per 2 μm). The HVC and LMAN inputs used in our simulation are assumed to arrive at one of the secondary dendrites shown in orange. Circuitry from the MSN to the LMAN neuron is simplified as a direct excitatory connection indicated by the dashed line. (B-E) Results obtained if the LMAN neuron synapses onto the dendritic shaft of the MSN. (B) Diagram of the secondary dendrite. LMAN input (red) arrives at the dendritic shaft in the middle. HVC inputs (blue) arrive at the spine heads. Simulated membrane voltages are sampled at the soma and the spine head that receives the HVC input, next to where the LMAN input arrives. (C) Examples of the membrane voltages of the soma (gray) and the recorded spine head that receives the HVC input (blue), when the HVC inputs are sub-threshold. The horizontal dashed line indicates the resting potential (-85 mV). (D) Calcium influx at the recorded spine head that receives the HVC input. The horizontal dashed line represents the threshold for synaptic plasticity. Vertical ticks at the bottom represent the timing of the HVC input (blue) and one example LMAN input (red). (E) The LMAN neuron’s firing probability throughout learning (calculated within a 10 ms window from 15 ms to 25 ms in a 50 ms simulation, see Methods). The shaded areas indicate the standard deviation across 10 repetitions. (F-I) Results obtained if the LMAN neuron synapses onto one spine head at the middle of the same secondary dendrite. (J) The LMAN neuron’s firing probability at trial 1000 as a function of the number of LMAN synapses onto spine heads that are evenly distributed along the same secondary dendrite. The vertical ticks indicate the standard deviation across 10 repetitions.

We next asked whether it is necessary for the LMAN neuron to synapse onto the MSN dendritic shaft, or whether the model still works if the LMAN neuron synapses onto the MSN spine heads (Fig. 3F). In this alternative configuration of the model, we observe that LMAN input onto the spine head produced little depolarization in neighboring spines, compared to LMAN input onto the shaft (Fig. 3G) because voltage attenuates strongly from spine head to shaft ^25,46,47^, resulting in insufficient calcium influx for plasticity (Fig. 3H). Consequently, the model failed to learn and the LMAN firing probability remained at the exploratory baseline (Fig. 3I).

However, the minimal depolarization of nearby spines permitted by restricting LMAN input onto a single spine head could be compensated for by having LMAN inputs onto multiple spine heads. We therefore quantified the amount of learning as a function of the number of spine heads with LMAN inputs. We found that at least 4 spine heads with LMAN synapses are required to drive learning, compared to a single shaft synapse (Fig. 3J), suggesting that LMAN input onto spine heads is approximately 4 times less efficient than LMAN input onto the shaft. The precise level of this efficiency improvement depended upon the input resistance of the MSN, but a similar many-fold improvement in efficiency held over the range of realistic input resistances (Fig. S3).

Connectomic analysis of brain circuitry has shown promise as a method to investigate synaptic plasticity ^48,49^. Using highly automated dense reconstruction, we have extended this approach to analyze the ultrastructure of synaptic interactions of inputs that carry computationally distinct learning-relevant signals. In the context of a cortical-basal ganglia circuit model that implements reinforcement learning, our data reveal the existence of a significant asymmetry between two types of cortico-striatal connections ^50,51^. The LMAN input resembles mammalian pyramidal-tract (PT) projections and carries an efference copy of a motor command signal, while the HVC input resembles mammalian intratelencephalic (IT) projections and carries a state (timing) signal. The predominance of HVC inputs onto dendritic spines and LMAN inputs onto shafts supports a biophysical model of basal ganglia learning in which action efference copy inputs modulate or gate plasticity of state inputs to MSNs. In this view, MSN spines detect and transiently maintain a memory of recently active action-state pairings. LMAN inputs onto dendritic spines, like the HVC inputs, may also constitute a state signal such that previous actions can be part of the state information for future actions.

Our results provide insight into a plausible biological implementation of node perturbation ^6^, a form of stochastic gradient descent learning that performs synaptic credit assignment without backpropagation. Conceptually, our model is a variant that could be called “remote node-perturbation”, since action feedback connections transmit network output variations back to the relevant synapses for credit assignment. We have identified a candidate ultrastructural motif for this mechanism: distinct excitatory synapse types interacting on a dendrite to inject variability, to correlate variability with reward signals, and to correctly assign credit to relevant synapses that potentiate improved performance. The generality of such a credit assignment strategy and whether it is manifested similarly in other brain circuits remains an interesting topic for future theoretical and empirical work.

## Material and Methods

### Sample preparation

Two adult male zebra finch (> 120 days post hatching, one obtained from the MPI of Ornithology, Seewiesen, Germany for the smaller j0126 data set; the second obtained from MIT breeding facilities for the larger j0251 data set) were transcardially perfused with cacodylate buffer (CB) at room temperature (RT) that contained 0.07 M sodium cacodylate (Serva, Germany), 0.14 M sucrose (Sigma-Aldrich), 2 mM CaCl_2_ (Sigma-Aldrich), 2% (1% for second finch, large data set sample) PFA (Serva) and 2% (1% for second finch) GA (Serva) to retain the extracellular space ^52^. Perfusion of the j0251 data set was performed by members of the Fee lab at MIT. We cut the brain sagittally into 200 µm (300 µm) thick slices on a vibratome (Leica VT1000S), found the slice with the largest cross section of Area X (identified by its round shape and location in the section) and excised a piece of about 300 x 300 µm (500 x 500 µm) centered in Area X. We next immersed the sample in a sequence of aqueous solutions in 2 ml Eppendorf tubes. First 2% OsO_4_ (Serva) reduced with 2.5 % potassium hexacyanoferrate(II) (Sigma-Aldrich) for 2 h at RT, followed by 1% thiocarbohydrazide (Sigma-Aldrich) at 58°C for 1h, then 2% OsO_4_ at RT for 2h, 1.5% uranyl acetate (Serva) in H_2_O at 53°C for 2h and 0.02M lead-aspartate (Sigma-Aldrich) at 53°C for 2h. The sample was rinsed three times with CB after the first OsO_4_ step and once with double-distilled water after each of the other staining steps. We next dehydrated the samples using chilled ethanol-water mixtures (70 %, 100 %, 100 %, 100 %, ethanol (Electron Microscopy Sciences), 10, 15, 10, 10 min) followed by propylene oxide (100%, 100%) (Sigma-Aldrich), infiltrated first with a 50/50 propylene oxide/epoxy mixture, then with 100% epoxy. The epoxy was 812-replacement (hard mixture, Serva). After infiltration, samples were cured (60°C for 48h), trimmed, gold-coated, and smoothed (Leica Ultracut microtome). Animal experiments for the smaller j0126 data set were approved by the Regierungspräsidium Karlsruhe and were carried out in accordance with the laws of the German federal government. Animal care and experiments for the larger j0251 data set were carried out in accordance with NIH guidelines, and reviewed and approved by the Massachusetts Institute of Technology Committee on Animal Care.

### SBEM data-set generation

The samples were imaged and sectioned in a field-emission scanning electron microscope (SEM, Zeiss UltraPlus) equipped with a custom diamond knife-based serial-blockface ultramicrotome ^27^. For the first sample (j0126), we used a landing energy of 1.6 kV, a beam current of 1 nA, a scan rate of 3.3 MHz, a lateral pixel size of 9 x 9 nm and a cutting thickness of 20 nm. The acquired data set (single image tile 10,240 x 10,240 pixels) was initially aligned by translation only, followed by elastic alignment (A. Pope, Google Research). The aligned stack contained 10,664 × 10,914 × 5701 voxels with 8 bit intensity values (∼664 GB). No contrast normalization was performed.

For the second sample (j0251) we used a landing energy of 2.0 kV, a beam current of 1 nA, a scan rate of 5.0 MHz, a lateral pixel size of 10 x 10 nm and a cutting thickness of 25 nm. This data set was acquired with multiple image tiles (4 x 4, 6,600 x 6,600 pixels) and first aligned using affine transformations ^53^, followed by elastic alignment (A. Pope, Google Research). The aligned stack contained 27,119 x 27,350 x 15,494 voxels with 8 bit intensity values (∼ 11 TB) and local contrast enhancement (CLAHE) was applied.

### Neuron reconstruction and proofreading

Neurite reconstruction was carried out by Flood Filling Network (FFN) segmentation of both data sets. Broadly, this was done in two stages: an initial over-segmentation (base segmentation) was created to ensure the near-absence of false mergers; next, segments in the base segmentation were then agglomerated into more complete units. Several steps were taken to improve oversegmentation in the initial stage (i.e. to further reduce the occurrence of false mergers), and to improve object continuity in the second stage (i.e. to reduce the occurrence of false splits). In addition to the base segmentation described in Januszewski et al. ^28^, we created an independent new oversegmentation that was then combined with the original base segmentation using oversegmentation consensus ^28^. The new oversegmentation started with the preexisting base segmentation as its initial state. We then used FFN inference with the following adjustments to create new neurite fragments :(a) we increased the “disconnected seed threshold” parameter from 0 to 0.15, which resulted in the assignment of previously unclaimed voxels in the interior of small-diameter axons (36.4B voxels filled); (b) we used a different snapshot of FFN weights (“checkpoint”) in areas with lower data quality (19.8B voxels filled); (c) we recreated objects adjacent to voxels classified as myelin by the tissue type classifier in an FFN inference run without the myelin mask (6.9B voxels filled); and (d) in areas adjacent to detected irregularities we recreated objects in an FFN inference run with FOV movement restriction with the section-to-section shift threshold relaxed from 4 to 6 voxels (1.8B voxels filled). The steps a)-d) were carried out sequentially, and the result of every step was combined with that of the previous step with oversegmentation consensus. To even further reduce the number of false mergers, we took the seed point for every segment in the original base segmentation, and used FFN inference at 2x reduced in-plane resolution to create a segmentation with no encumbrance by any other object (note that in this process the base-set objects can shrink as well as grow). The results were upsampled and combined with the base segmentation, again through oversegmentation consensus.

We then ran FFN agglomeration as described previously ^28^. For every decision point involving at least one segment added in (a)-(d) (see paragraph above), we ran FFN agglomeration a second time, this time with inference settings matching the conditions (a)-(d) under which the segment was created. To detect remaining splits we skeletonized the agglomerated objects using TEASAR ^54^. For each skeleton node with only one adjacent edge we determined whether it was a true neurite endpoint by running FFN segmentation within an empty (no pre-existing segments) (201,201,101)-voxel subvolume centered on the seed placed at the skeleton node. This was done with both the main FFN checkpoint and the one used in condition (b) above. We considered any base segments to be “recovered” if at least 60% of their voxels were overlapped by the predicted object map (POM) generated by the FFN in this procedure, and thresholded at 0.5. We then merged segments (A, B) in case of symmetric recovery, i.e. when the subvolume segmentation described above, seeded from a node located in A recovered B, and vice versa. Finally, we identified orphan fragments (defined as not reaching the surface of the data set) and iteratively (6 times) connected them to the partner fragment with the highest Jaccard score as computed in FFN agglomeration for the fragment pair^28^. Only pairs for which at most 15% of the voxels of one of the objects changed POM values from >0.8 to <0.5 during FFN agglomeration inference were considered in this step.

The number of orphaned neurite fragments in the small data set was further reduced by manually inspecting their ends. If a split error was detected, a skeleton tracing was initiated. The annotator was provided with a seed location and an initial tracing direction but not the actual segmentation. Tracing was terminated when the annotator decided that a true neurite ending had been reached or when the newly traced skeleton reached another fragment with > 10 µm skeleton length. The thus created connectors (n=36,154 in 911 hours of tracing) were then used to combine the appropriate fragments. This reduced the number of fragments from 231,207 to 212,656. 13 of those contained at least one erroneous merger, identified by the presence of at least two somata. All connectors contained in those (n=434) were then removed, resulting in 213,076 fragments. This proofreading step was performed using a Python plugin for KNOSSOS (www.knossostool.org).

To further reduce the number of false mergers in the small data set, one of us (MJ) inspected 36,653 classified neurites using Neuroglancer, identified merge errors by visual evaluation of object shape plausibility, and manually removed agglomeration graph edges causing those mergers. In total, less than 10 hours were spent for this additional proofreading step, which resulted in the removal of 846 edges and the elimination of 840 merge errors.

Unless otherwise mentioned, manual proofreading was performed by student assistants or outsourced (ariadne.ai ag).

### Synapse detection

In a first step, mitochondria, vesicle clouds, synaptic junctions and synapse type were predicted using two convolutional neural networks (CNN) in the j0126 data set described in detail in ^29^. The same classes were predicted in the j0251 data set using a single CNN, with further processing described in detail in ^31^ and in the companion paper by Rother et al., 2025. The resulting synaptic junction predictions were turned into candidate synapse objects through connected components.

In a second step, every candidate synapse object was assigned a probability by a random forest (RF) classifier trained on manually labeled candidate synapse objects (j0126 data set: n=436; non-synaptic: 238, synaptic: 198). The synaptic area of identified synapses was estimated as its surface mesh area (which includes pre- and post-synaptic areas) divided by two.

### Cellular compartment classification

We used cellular morphology neural networks (CMN) ^30^ for the j0126 data set to recognize and differentiate cellular substructures in the small data set. The two models, one for spines ^30^ and one for axons (axon, terminal bouton, *en-passant* bouton, dendrite, soma; trained on 45 reconstructions; window size of 40.96 µm x 20.48 µm x 20.48 µm and 1024 by 512 pixels; 3 projection windows per location; rendering locations were the set of vertices downsampled by a third of the maximum window size), classified the cell surface based on 2D-projections of the neurite and its mapped organelles (mitochondria, vesicle clouds) and synaptic junctions. Surface labels were mapped to the synapse directly by k-nearest-neighbors in case of spines (k=50). In the axon (dendrite, soma and bouton) case the surface labels were first mapped to the skeleton nodes by k-nearest-neighbors (k=50) and then averaged using a sliding window approach, which collected the labels of skeleton nodes within 10 µm traversal length along the skeleton in every direction starting from the source node. The axon (dendrite, soma, bouton) label of each synaptic partner was taken from the respective skeleton node closest to the synapse.

For j0251, we used a point-cloud based morphology classifier for the neuronal compartment classification, as described in ^31^ and in detail in the companion paper Rother et al., 2025.

### Cell-type classification

The cell types in the j0126 data set of the remaining FFN reconstructions were then identified with the same approach, with a CMN trained on manually labeled neurites (JK), 30 per class, in total n=240 (HVC, LMAN, MSN, pallidal, subthalamic nucleus like, dopaminergic, cholinergic, interneuron; the dopaminergic and cholinergic classes were combined for the classification used in this manuscript). We modified the CMN from ^30^ by extending the latent representation returned by the last convolution layer with the synaptic area ratio between symmetric and all synapses in the neurite before it was passed to the first fully connected layer. The performance was assessed using ten-fold cross-validation ^55^.

The cell types in the large data set were classified as described in ^31^ through a point-cloud based approach, with an extended set of ground truth cell types, as the large data set allowed a far more comprehensive manual classification (see companion paper Rother et al., 2025).

### Synapse analyses of HVC and LMAN axons

Synaptic size (contact area) was computed as described in the ‘Synapse detection’ section of the methods.

Synapse sizes were log-10 transformed and the distribution was fit to a mixture of two truncated (lower boundary of 0.01 µm^2^, no upper boundary) normal distributions (Python scipy.stats.truncnorm) by minimizing the negative log-likelihood function using Sequential Least Squares Programming (Python scipy.optimize.fmin_slsqp).

We calculated the fraction of total synaptic area that HVC and LMAN axons dedicate to the different postsynaptic compartments of MSNs. This analysis was restricted to synapses from automatically classified axons (SyConn celltype_certainty > 0.8; synapse probability > 0.6) onto well-reconstructed MSNs, which required at least 200 µm of dendritic, 50 µm of axonic, and 5 µm of somatic skeleton length. Postsynaptic compartments, such as spine heads, necks, dendritic shafts, or somas, were identified using a morphology classifier ^30,31^. For each presynaptic axon type, the area fraction was then computed by dividing the total synaptic area onto one compartment class by the total synaptic area onto all MSN compartments. For example, the data set-wide spine fraction for HVC onto MSNs was computed as follows:

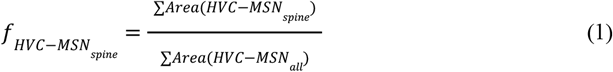

For the manually verified ground-truth axons, the synaptic area fraction was calculated by dividing the total area onto spine heads by the sum of the areas on both spine heads and dendritic shafts. To assess if targeting preferences were consistent across individual neurites, we also analyzed distributions on a per-axon (Fig. 2C) and per-dendrite (Fig. 2D) basis. The axon-centric analysis included both the manually verified ground-truth population and the larger pool of automatically classified axons (celltype_certainty > 0.8), filtering for individual axons with at least 50 synapses and a skeleton length of 50 µm or more. The dendrite-centric analysis was similarly restricted to MSN dendrites with at least 50 synapses, and their input fractions were calculated using only presynaptic partners that met the same 50 µm axon length criterion.

We performed a similar analysis on the smaller dataset j0126 with slightly adjusted filtering criteria to account for the smaller data set size: a synapse probability of >0.5, and a minimum axon length and synapse count of 10 µm and 10, respectively. We again calculated synaptic area fractions onto MSN compartments, analyzed the per-neurite targeting distributions, and characterized the synapse size distributions with truncated normal mixtures to confirm the original findings.

To quantify targeting preference, we performed an enrichment analysis based on the synaptic contact area for auto-classified axons. We first computed the conditional probability of a synapse targeting a specific postsynaptic compartment given its origin (e.g., p(shaft|LMAN)) and the corresponding marginal probability across all synapses (e.g., p(shaft)). The enrichment factor was then calculated as the ratio of the conditional to the marginal probability, indicating how much more likely a given axon class is to target a specific compartment compared to the overall average.

### Computational model of reinforcement learning in Area X

The neural circuit model of reinforcement learning in Area X is based on a reconstructed MSN that receives the HVC and LMAN input, as well as a reinforcing dopaminergic signal, and that drives the LMAN neuron. The LMAN neuron is modeled as a conductance-based, leaky integrate-and-fire neuron since we only consider its spike timing and firing probability. The HVC input is modeled simply as a train of spikes driving song production at a given note. We first describe the neural circuit model in general and then describe the model details of the MSN and LMAN neuron. Computer code and data files containing model parameters are provided in the Supplementary Files.

#### The neural circuit model

Each trial of learning lasts 50 ms, representing 5 notes of the birdsong. We consider each discrete 10 ms window as one note (0-10 ms, 10-20 ms, etc.). Simulations were performed in Python with a simulation time step of 0.05 ms, using the NEURON library for simulations of the MSN.

The circuit connectivity and synaptic learning rule are as follows: The MSN provides excitatory synaptic input to the LMAN neuron with a 10 ms delay. This delay approximates the circuit delay associated with the polysynaptic circuit from the MSN, through the pallidal-like and thalamic neurons, to the LMAN neuron. The LMAN neuron, with 1 ms delay, synapses onto either the dendritic shaft (Fig. 3B-E) or one spine head (Fig. 3F-I) located in the middle of an arbitrarily selected secondary dendrite of the MSN, or multiple spine heads that are evenly distributed along the same secondary dendrite (Fig. 3J, Fig. S3). The MSN also receives HVC input at all spine heads of the selected secondary dendrite (except for the spine head or heads receiving LMAN input). The HVC input is modeled as a burst of four identical spikes within a 10 ms window at the beginning of the 50 ms simulation (at 2, 4, 6, and 8 ms relative to simulation start), representing one note of the birdsong. The LMAN-MSN synapse is assumed to be non-plastic, while the HVC-MSN synapses are plastic. The synaptic weight of each HVC-MSN synapse in one trial of singing is updated according to the following learning rule, which is applied at each discrete 10 ms window:

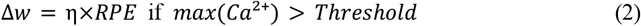

where Δ*w* is the change of synaptic weight, η is the learning rate, and *RPE* is the reward prediction error calculated for each note (see below). *max*(*Ca*^2+^) is the maximum amplitude of the calcium current mediated by the NMDA receptor. *Threshold* is the threshold for synaptic plasticity. Parameter values are described in detail in the section *The MSN model* below.

Reward-based learning is governed by the timing of the LMAN spikes. For simplicity, we consider only a single LMAN neuron. We assume that this neuron generates exploratory spikes that improve the song performance. The goal of learning is to increase the firing probability of this LMAN neuron within a window between 15 ms and 25 ms relative to simulation start, which, given the HVC to LMAN circuit delay, approximately corresponds to reinforcing the HVC input that arrived within the first 10 ms window of the simulation. A reward (*R*) is defined for each discrete 10 ms note as a function of the spike timing of the LMAN neuron. The reward takes a maximum value of 1 when the LMAN neuron generates a spike at 20 ms relative to simulation start, and falls off according to a Gaussian function with a standard deviation of 10 ms. If the LMAN neuron generates more than one spike within a note, one of the spikes is randomly selected to calculate the reward. A reward prediction (*RP*) is maintained for each discrete 10 ms note and updated at each trial according to the following update rule:

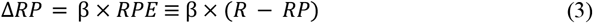

where β = 0. 1 is the update speed.

#### The LMAN neuron model

The LMAN neuron is modeled as a conductance-based, leaky integrate-and-fire neuron. The subthreshold dynamics follows:

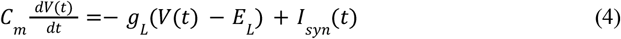

where *V*(*t*) is the membrane voltage, *C*_*m*_ = 0. 5 nF is the membrane capacitance, *g* _*L*_ = 25 nS is the leak conductance, and *E* _*L*_ =− 65 mV is the resting potential. *I* _*syn*_ (*t*) represents the total synaptic input, consisting of the excitatory MSN input, an excitatory background Poisson spike train, and an inhibitory background Poisson spike train:

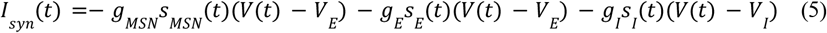

Where *g* _*MSN*_ = 20 nS, *g* _*E*_ = 5 nS, and *g* _*I*_ = 5 nS are the maximal conductances of the excitatory MSN-LMAN synaptic connection, the excitatory background input, and the inhibitory background input, respectively. *V* _*E*_ = 0 mV is the reversal potential of the excitatory input, modeling AMPA receptors, and *V* _*I*_ =− 70 mV is the reversal potential of the inhibitory input, modeling GABA receptors. *s*(*t*) represents the time course of activation of the corresponding input, modeled as a double exponential with fast kinetics mimicking those of fast AMPA ^56^ and GABA ^57^ receptors:

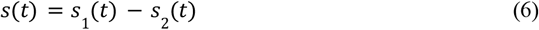

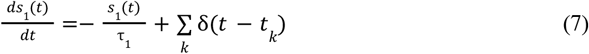

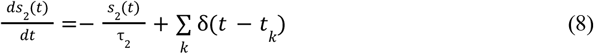

Where τ_1_ = 1. 0 ms and τ_2_ = 0. 1 ms. Here, δ is a delta function such that *s*_1_ (*t*) and *s*_2_ (*t*) increases by 1 when there is a presynaptic spike at time *t* = *t*_*k*_. The background Poisson input firing rates were set to 2.6 kHz for excitatory background and 3.0 kHz for inhibitory background. These values were chosen to elicit a baseline exploratory firing probability near 0.2 within a 10 ms note, approximating the spontaneous firing rate of LMAN neurons during singing without MSN input ^58,59^. The spiking threshold *V*_*thr*_ was set to − 50 mV. The reset voltage, *V*_*reset*_ =− 53 mV, was set to be slightly below *V*_*thr*_, modeling the tendency of the voltage of cortical neurons to hover near threshold. The refractory period was set to 1 ms.

#### The MSN model

The MSN was constructed with morphology based on the 3D reconstruction shown in Fig. 3A and ion channel dynamics described in detail below. Simulations of the MSN were performed using the NEURON library in Python.

##### Morphology

The soma of the MSN has a length of 10 μm and a diameter of 10 μm. There are 5 primary dendrites (labeled from 1 to 5) with morphology specified in Table 1.

**Table 1:**
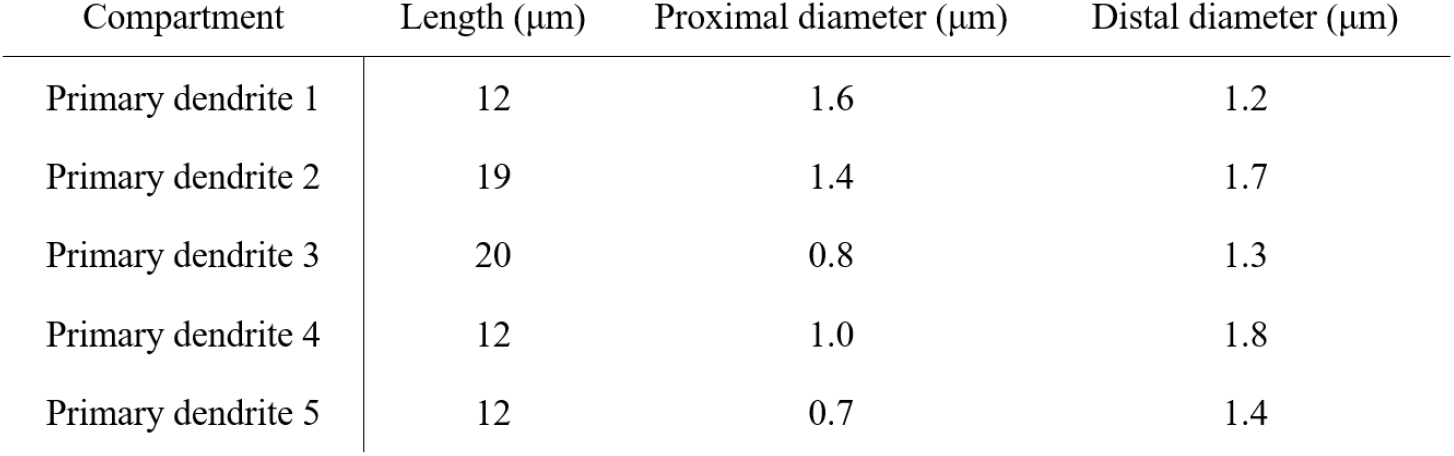
Morphology of the primary dendrites.

| Compartment | Length ($\mu\text{m}$) | Proximal diameter ($\mu\text{m}$) | Distal diameter ($\mu\text{m}$) |
| --- | --- | --- | --- |
| Primary dendrite 1 | 12 | 1.6 | 1.2 |
| Primary dendrite 2 | 19 | 1.4 | 1.7 |
| Primary dendrite 3 | 20 | 0.8 | 1.3 |
| Primary dendrite 4 | 12 | 1.0 | 1.8 |
| Primary dendrite 5 | 12 | 0.7 | 1.4 |

The diameter of each dendritic compartment in the model linearly changes from the proximal diameter to the distal diameter. The actual diameters of the dendrites in the 3D reconstruction vary irregularly at different locations. The parameters listed above correspond to manual measurements of dendritic diameters at the base of each dendrite and at the branching points. Primary dendrite 1 is attached to one end of the soma. Primary dendrites 2 and 3 are attached to the other end of the soma. Primary dendrites 4 and 5 are attached to the middle of the soma.

There are 2 to 4 secondary dendrites on each primary dendrite (labeled as 1-1, 1-2, 1-3, 2-1, etc.), with geometries specified in Table 2. Secondary dendrite 1-1 is attached to the 7 μm location (from proximal to distal) on primary dendrite 1. Secondary dendrite 2-1 is attached to the 7 μm location on primary dendrite 2. Secondary dendrite 2-2 is attached to the 10 μm location on primary dendrite 2. Secondary dendrite 3-1 is attached to the 11 μm location on primary dendrite 3. All other secondary dendrites are attached to the distal end of the corresponding primary dendrites.

**Table 2:**
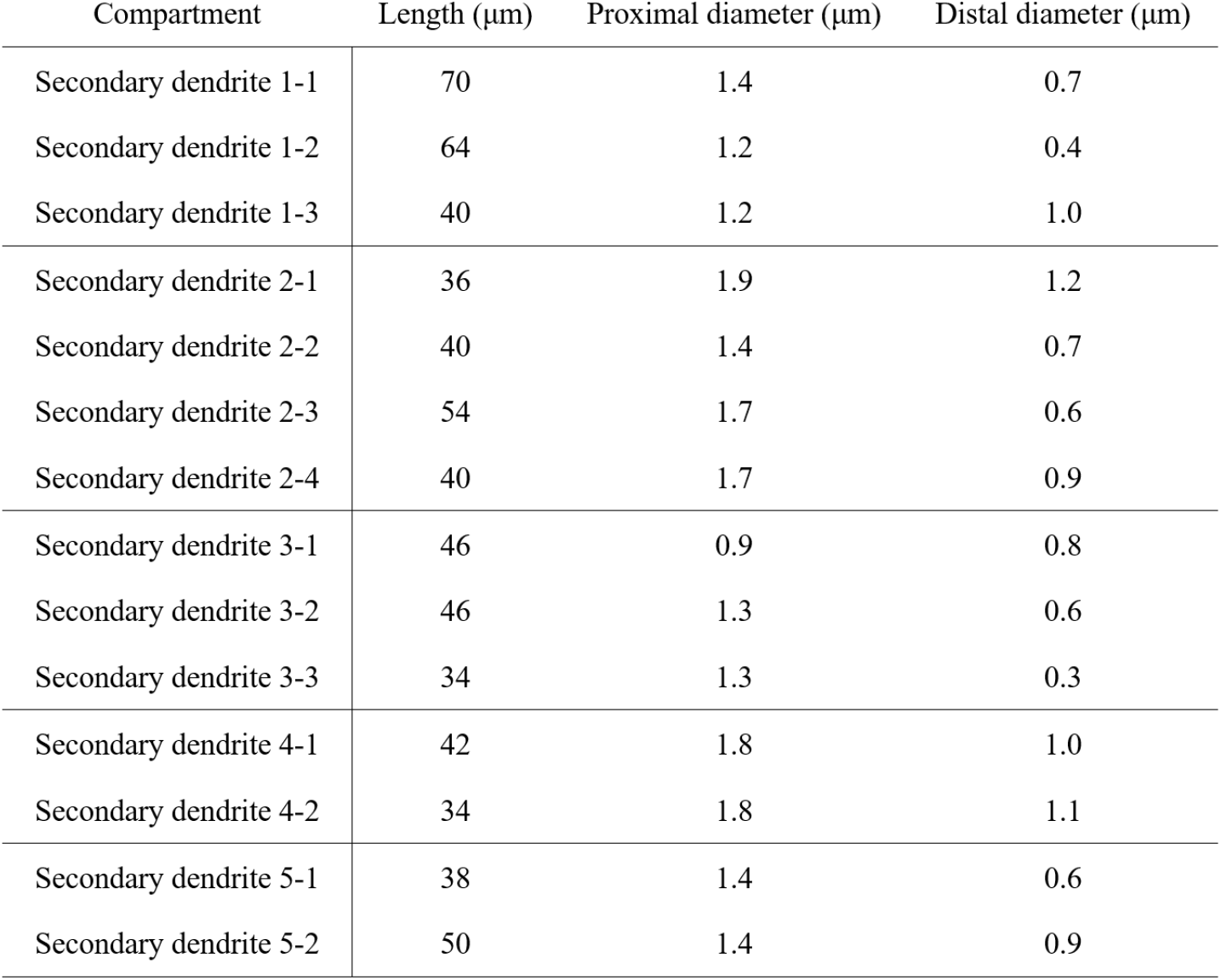
Morphology of the secondary dendrites.

The selected secondary dendrite that receives HVC and LMAN inputs is 1-2. For the simulations in Fig. 3 and Fig. S3, spines are evenly distributed on all secondary dendrites with a density of one spine per 2 μm, with spine morphology as follows ^60^: head length and diameter are both 0.5 μm; neck length is 0.7 μm; neck width is 0.15 μm. For the simulations of Fig. S4, see “Model with detailed spine location and morphology” below.

##### Voltage and ion channel dynamics

Each spine head, spine neck, and each 1 μm of the soma and the dendrites is modeled as one isopotential segment. The membrane voltage of each segment is modeled as

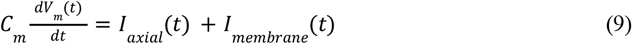

where *C* _*m*_ = 1.0 μF/cm^2^ is the membrane capacitance per unit area. The current received by each segment consists of axial currents, *I*_*axial*_ (*t*), from nearby segments, and membrane currents, *I*_*membrane*_(*t*), conducted by membrane ion channels including those activated by synaptic inputs. The axial resistivity of all segments except for the spine necks is 100 Ω·cm. The axial resistivity of the spine necks is 1000 Ω·cm to approximate the relatively high spine neck resistance measured in experiments ^25^. The membranes of all segments (including soma, primary dendrites, secondary dendrites, and spines) are embedded with inward rectifying potassium channels (Kir). The membranes of the soma and the primary dendrites are also embedded with 4-AP resistant persistent potassium channels (Krp), fast A-type potassium channels (Kaf), and fast sodium channels (Naf). Channel dynamics are adapted from ^61^ and described below. We elevated the *q* factor from 0.5 to 1 for the Kir channels, and from 3 to 5 for the Krp, Kaf, and Naf channels, to account for the higher body temperature in songbird.

The sodium and potassium currents were modeled in the Hodgkin-Huxley formalism by the equation

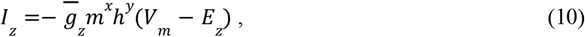

where the subscript *z* denotes each channel type except for the Krp channel, which is described as partially inactivating ^62^:

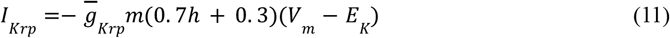

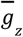 is the maximal conductance per unit area (see Table 3). *E*_*z*_ is the reversal potential for each ion species: *E*_*Na*_ = 50 mV for sodium and *E*_*K*_ =− 85 mV for potassium. Values of the exponent *x* and *y* depend on the specific channel type (see Table 3). *m* and *h* represent the activation and inactivation state of the channel, respectively, and are calculated according to:

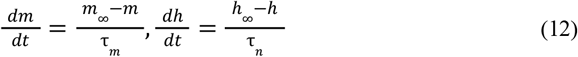

where *m*_∞_, *h*_∞_, τ_*m*_, and τ_*h*_ are functions of the membrane voltage, *V*_*m*_, and represent the steady state activation curves and time constants for *m* and *h. m*_∞_ and *h*_∞_ are described by equations of the form (they have the same form and thus we only write *m*_∞_):

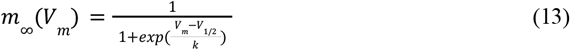

where *V*_1/2_ and *k* represent the half-activation voltages and slopes of the steady-state Boltzmann fits to *m*_∞_ and *h*_∞_. The tabulated time constants ^61^ are provided in the Supplementary Files. Parameters of the channel dynamics are summarized in Table 3.

**Table 3:** Parameters used for membrane ion channels in the model.

| Channel type | $\bar{g}$ (S/cm<sup>2</sup>) | HH form | | $V_{1/2}$(mV) | $k$(mV) | $\tau$(ms) |
| --- | --- | --- | --- | --- | --- | --- |
| Kir | Varied | $m$ | $m$ | -82.0 | 13.0 | Tabulated |
| Krp | 1e-2 | $m(0.7h+0.3)$ | $m$ | -13.5 | -11.8 | Tabulated |
| | | | $h$ | -54.7 | 18.6 | Tabulated |
| Kaf | 1.5 | $m^2h$ | $m$ | -10.0 | -17.7 | Tabulated |
| | | | $h$ | -75.6 | 10.0 | 4.67 |
| Naf | 3.0 | $m^3h$ | $m$ | -23.9 | -11.8 | Tabulated |
| | | | $h$ | -62.9 | 10.7 | Tabulated |

The values of the maximal conductances of the Kir channels were chosen to achieve input resistances consistent with those measured experimentally in striatal medium spiny neurons. Specifically, three different maximal conductances per unit area were used to achieve different input resistances measured at the soma at the resting potential of -85 mV (Fig. S3): 1e-3 S/cm^2^ leads to around 60 MΩ; 5e-4 S/cm^2^ leads to around 100 MΩ; and 2.5e-4 S/cm^2^ leads to around 170 MΩ. These input resistances fall within the experimentally observed range for MSNs around this voltage in slice recordings, which spans from about 50 MΩ ^63^ to 200 MΩ ^64^, with intermediate values reported in other studies ^61,65^. Notably, much higher input resistances have also been reported: around 380 MΩ ^66^ and 670 MΩ ^34^. However, the “input resistances” in these two studies were defined as the maximum slope of the measured current-voltage (I-V) curve at membrane potentials below -50 mV, rather than the input resistance measured at rest. Examination of the I-V curves in these studies around the resting potential suggest an input resistance around 100 MΩ, consistent with the other studies. The reason for the higher input resistances at depolarized potentials is that the Kir channels, which dominate the resting potentials of the MSNs, inactivate when the cell is depolarized ^67^. In our model, if we remove the fast sodium (Naf) and potassium (Kaf) channels that predominantly contribute to action potentials, the input resistance of the model neuron similarly increases from around 100 MΩ at rest to maximally around 700 MΩ when depolarized, demonstrating that Kir inactivation during depolarization accounts for the apparent change in input resistance.

The LMAN synaptic input is assumed to be mediated by AMPA receptors with the same dynamics described above:

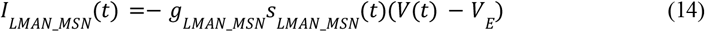

The maximal conductance of the LMAN-MSN synaptic input, *g*_*LMAN*_*MSN*_, is fixed at 20 nS. To make the comparison between the model performance when the LMAN input synapses onto the dendritic shaft (Fig. 3B-E) versus spine head (Fig. 3F-J), the same maximal conductance is used in both configurations. In the latter case (Fig. 3F-J), the LMAN spine input produces a depolarization near saturation in the spine head (close to the excitatory reversal potential of 0 mV, not shown in Fig. 3 for simplicity). This depolarization attenuates strongly when it propagates through the spine neck to the dendritic shaft (Fig. 3G).

The HVC synaptic input is assumed to be mediated by both AMPA and NMDA receptors. The AMPA current is modeled with the same dynamics described above:

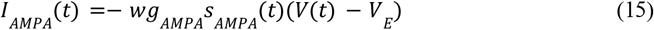

where *g*_*AMPA*_ = 0. 05 nS. To describe the plasticity occurring across the time course of learning, we define a synaptic weight parameter *w* (see Equation 2) that scales the amplitude of the AMPA conductance. The synaptic weight is initialized to *w* = 1. The learning rate is η = 1. The NMDA current has a voltage dependence that is a function of the extracellular magnesium concentration ^68^, and is modeled as:

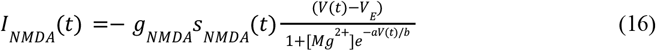

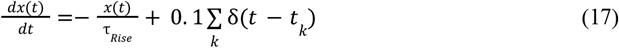

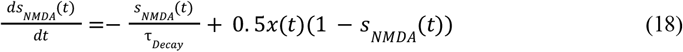

Where *g*_*NMDA*_ = 0. 3 nS (for simulations with an input resistance of 60 MΩ and 100 MΩ) or *g*_*NMDA*_= 0. 2 nS (for simulations with an input resistance of 170 MΩ). [*Mg*^2+^] = 1 represents the extracellular magnesium concentration. *a* = 0. 062, *b* = 3. 57, τ_*Rise*_ = 15 ms, and τ _*Decay*_ = 30 ms.

The calcium influx is mediated by the NMDA receptors. We consider the conductance of calcium to be 10 percent of the total NMDA conductance and model the calcium influx by:

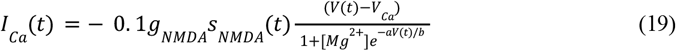

Where *V*_*Ca*_ = 125 *mV* is the reversal potential of the calcium conductance. The threshold for plasticity is *Threshold* = 0. 8 pA (for simulations with an input resistance of 60 MΩ and 100 MΩ) or *Threshold* = 0. 6 pA (for simulations with an input resistance of 170 MΩ).

##### Model with detailed spine location and morphology

As noted above, for the simulations shown in Fig. 3 and Fig. S3, we used a uniform spatial distribution of the synapses onto each MSN secondary dendrite and each spine had the same morphology. To verify that this simplification did not change the main results, in Fig. S4 we simulated a model that used the measured spatial locations of the synapses onto the modeled MSN and the measured length and diameter of each individual spine. In this model, there are 25 HVC synapses onto the spine heads distributed along the selected secondary dendrite and one LMAN synapse onto the dendritic shaft at 28.85% of the total length of the selected secondary dendrite from proximal to distal. The fourth HVC synapse (from proximal to distal) is closest to the LMAN synapse and is recorded for plotting. To make the comparison between the model performance when the LMAN input synapses onto the dendritic shaft (Fig. S4A-C) versus spine head (Fig. S4D-F), the LMAN synapse onto the dendritic shaft is replaced with one onto the head of a spine with the same morphology as used in the model of Fig. 3 and Fig. S3. For this simulation,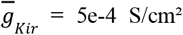 leading to an approximately 100 MΩ input resistance measured at the soma at the resting potential of -85 mV, *g*_*NMDA*_ = 0. 2 nS, *g*_*LMAN*_*MSN*_ = 30 nS, and the threshold for plasticity is *Threshold =* 0.5 pA. The other parameters are the same as described above.

## Acknowledgements

We thank Lorenz Hüdepohl and Christian Guggenberger for their support at the MPCDF in Garching, and Manfred Gahr for providing zebra finches. We thank Ila Fiete, Jesse Goldberg and John Shlens for comments on an earlier version of the manuscript. We would also like to thank the KNOSSOS team for programming support, and Marius Killinger and Martin Drawitsch for help with the ELEKTRONN library. We also thank Julia Kuhl for her assistance with figure preparation. Our thanks go to Sven Dorkenwald, Hashir Ahmad and Andrei Mancu for help with SyConn processing and development. We further thank Riccardo Morbio, Delta Schick, and Laura Werner for the manual verification of synapses used in the RFC ground truth and data annotation in general. We especially thank Riccardo Morbio for additionally evaluating RFC predictions on synapses and reviewing cells for mergers and autapses. We are also grateful to the annotators who contributed to the synapse and mitochondria ground truth: Julian Hendricks, Delta Schick, Riccardo Morbio, Katyayni Ganesan, Deniz Üreyener, Laura Werner, Merve Cetiner, Gizem Karabiyik, Yona Perstat, Ata Kan, Dominik Melzer, Akshaya Rajan, and Maria Nikitina. Finally, we thank Delta Schick and Julian Hendricks for reviewing the annotations. This work was funded by the Max Planck Society, Google Research, grant NIH RF1 MH117809 and UKRI.

## Code availability

Data was prepared using the SyConn2 pipeline ^31^, [SyConn repository](https://github.com/StructuralNeurobiologyLab/SyConn.git) ^31^. All analysis code will be made available upon final publication. The code of the biophysical model is available in a [GitHub repository](https://github.com/goldman-lab/Biophysical-model-of-credit-assignment).

## Data availability

The datasets are available on SyConn Web (syconn.esc.mpcdf.mpg.de) under the dataset ‘j0126’ and ‘j0251’.

## Ethics declarations

Competing interests: JK holds shares of ariadne.ai ag.

## Author contributions

JK collected the j0251 and j0126 volume EM data set. WD supervised data acquisition and developed instrumentation with contributions by JK. JK and AR annotated training data and coordinated data annotation with contributions from MJ. JK and MJ processed and aligned the volume EM data. PS, AR and JK coordinated and performed the connectome extraction with SyConn v2, based on EM segmentations by MJ. JK analysed the connectome and prepared figures, with contributions from AR. YW implemented the biophysical model, supervised by MG, with contributions from MSF and JK. JK, YW, MG, MJ, VJ, WD and MSF wrote the manuscript. JK, WD, YW, MG, VJ and MSF conceptualized the study. All authors reviewed the manuscript.

## References

1. Rumelhart, D. E., Hinton, G. E. & Williams, R. J. Learning representations by back-propagating errors. Nature vol. 323 533–536 Preprint at 10.1038/323533a0 (1986).

2. Guerguiev, J., Lillicrap, T. P. & Richards, B. A. Towards deep learning with segregated dendrites. Elife 6, (2017).

3. Lillicrap, T. P., Cownden, D., Tweed, D. B. & Akerman, C. J. Random synaptic feedback weights support error backpropagation for deep learning. Nat. Commun. 7, 13276 (2016).

4. Bartunov, S. et al. Assessing the Scalability of Biologically-Motivated Deep Learning Algorithms and Architectures. in Advances in Neural Information Processing Systems 31 (eds. Bengio, S.et al.) 9368–9378 (Curran Associates, Inc., 2018).

5. Williams, R. J. Simple Statistical Gradient-Following Algorithms for Connectionist Reinforcement Learning. Reinforcement Learning 5–32 Preprint at 10.1007/978-1-4615-3618-5_2 (1992).

6. Fiete, I. R. & Seung, H. S. Gradient learning in spiking neural networks by dynamic perturbation of conductances. Phys. Rev. Lett. 97, 048104 (2006).

7. Thorndike, E. L. Animal intelligence: An experimental study of the associative processes in animals. Preprint at 10.1037/10780-000 (1898).

8. Sutton, R. S., Barto, A. G., Co-Director Autonomous Learning Laboratory Andrew G Barto & Bach, F. Reinforcement Learning: An Introduction. (MIT Press, 1998).

9. Doya, K. What are the computations of the cerebellum, the basal ganglia and the cerebral cortex? Neural Netw. 12, 961–974 (1999).

10. Brainard, M. S. & Doupe, A. J. Translating birdsong: songbirds as a model for basic and applied medical research. Annu. Rev. Neurosci. 36, 489–517 (2013).

11. Gale, S. D. & Perkel, D. J. Anatomy of a songbird basal ganglia circuit essential for vocal learning and plasticity. J. Chem. Neuroanat. 39, 124–131 (2010).

12. Olveczky, B. P., Andalman, A. S. & Fee, M. S. Vocal experimentation in the juvenile songbird requires a basal ganglia circuit. PLoS Biol 3, e153 (2005).

13. Kao, M. H., Doupe, A. J. & Brainard, M. S. Contributions of an avian basal ganglia–forebrain circuit to real-time modulation of song. Nature vol. 433 638–643 Preprint at 10.1038/nature03127 (2005).

14. Hahnloser, R. H. R., Kozhevnikov, A. A. & Fee, M. S. An ultra-sparse code underlies the generation of neural sequences in a songbird. Nature 419, 65–70 (2002).

15. Gadagkar, V. et al. Dopamine neurons encode performance error in singing birds. Science 354, 1278–1282 (2016).

16. Fee, M. S. & Goldberg, J. H. A hypothesis for basal ganglia-dependent reinforcement learning in the songbird. Neuroscience 198, 152–170 (2011).

17. Yagishita, S. et al. A critical time window for dopamine actions on the structural plasticity of dendritic spines. Science 345, 1616–1620 (2014).

18. Fee, M. S. Oculomotor learning revisited: a model of reinforcement learning in the basal ganglia incorporating an efference copy of motor actions. Frontiers in Neural Circuits 6, (2012).

19. Fee, M. S. The role of efference copy in striatal learning. Curr. Opin. Neurobiol. 25, 194–200 (2014).

20. Redondo, R. L. & Morris, R. G. M. Making memories last: the synaptic tagging and capture hypothesis. Nat. Rev. Neurosci. 12, 17–30 (2011).

21. Farries, M. A. & Fairhall, A. L. Reinforcement learning with modulated spike timing dependent synaptic plasticity. J. Neurophysiol. 98, 3648–3665 (2007).

22. Frémaux, N. & Gerstner, W. Neuromodulated Spike-Timing-Dependent Plasticity, and Theory of Three-Factor Learning Rules. Front. Neural Circuits 9, 85 (2015).

23. Svoboda, K., Tank, D. W. & Denk, W. Direct measurement of coupling between dendritic spines and shafts. Science 272, 716–719 (1996).

24. Yuste, R. & Denk, W. Dendritic spines as basic functional units of neuronal integration. Nature 375, 682–684 (1995).

25. Harnett, M. T., Makara, J. K., Spruston, N., Kath, W. L. & Magee, J. C. Synaptic amplification by dendritic spines enhances input cooperativity. Nature 491, 599–602 (2012).

26. Yuste, R. & Bonhoeffer, T. Morphological Changes in Dendritic Spines Associated with Long-Term Synaptic Plasticity. Annual Review of Neuroscience vol. 24 1071–1089 Preprint at 10.1146/annurev.neuro.24.1.1071 (2001).

27. Denk, W. & Horstmann, H. Serial block-face scanning electron microscopy to reconstruct three-dimensional tissue nanostructure. PLoS Biol. 2, e329 (2004).

28. Januszewski, M. et al. High-precision automated reconstruction of neurons with flood-filling networks. Nat. Methods 15, 605–610 (2018).

29. Dorkenwald, S. et al. Automated synaptic connectivity inference for volume electron microscopy. Nat. Methods 14, 435–442 (2017).

30. Schubert, P. J., Dorkenwald, S., Januszewski, M., Jain, V. & Kornfeld, J. Learning cellular morphology with neural networks. Nat. Commun. 10, 2736 (2019).

31. Schubert, P. J. et al. SyConn2: dense synaptic connectivity inference for volume electron microscopy. Nat Methods 19, 1367–1370 (2022).

32. Fortune, E. S. & Margoliash, D. Parallel pathways and convergence onto HVc and adjacent neostriatum of adult zebra finches (Taeniopygia guttata). The Journal of Comparative Neurology vol. 360 413–441 Preprint at 10.1002/cne.903600305 (1995).

33. Vates, G. E. & Nottebohm, F. Feedback circuitry within a song-learning pathway. Proc. Natl. Acad. Sci. U. S. A. 92, 5139–5143 (1995).

34. Farries, M. A. & Perkel, D. J. A telencephalic nucleus essential for song learning contains neurons with physiological characteristics of both striatum and globus pallidus. J. Neurosci. 22, 3776–3787 (2002).

35. Bottjer, S. W. & Sengelaub, D. R. Cell death during development of a forebrain nucleus involved with vocal learning in zebra finches. J. Neurobiol. 20, 609–618 (1989).

36. Walton, C., Pariser, E. & Nottebohm, F. The Zebra Finch Paradox: Song Is Little Changed, But Number of Neurons Doubles. Journal of Neuroscience vol. 32 761–774 Preprint at 10.1523/jneurosci.3434-11.2012 (2012).

37. Goldberg, J. H. & Fee, M. S. Singing-related neural activity distinguishes four classes of putative striatal neurons in the songbird basal ganglia. J. Neurophysiol. 103, 2002–2014 (2010).

38. Dorkenwald, S. et al. Binary and analog variation of synapses between cortical pyramidal neurons. bioRxiv 2019.12.29.890319 (2019) doi:10.1101/2019.12.29.890319.

39. Fee, M. S. & Goldberg, J. H. A hypothesis for basal ganglia-dependent reinforcement learning in the songbird. Neuroscience 198, 152–170 (2011).

40. Leblois, A., Bodor, A. L., Person, A. L. & Perkel, D. J. Millisecond timescale disinhibition mediates fast information transmission through an avian basal ganglia loop. J Neurosci 29, 15420–15433 (2009).

41. Hamaguchi, K. & Mooney, R. Recurrent interactions between the input and output of a songbird cortico-basal ganglia pathway are implicated in vocal sequence variability. J Neurosci 32, 11671–11687 (2012).

42. Shindou, T., Ochi-Shindou, M. & Wickens, J. R. A Ca(2+) threshold for induction of spike-timing-dependent depression in the mouse striatum. J Neurosci 31, 13015–13022 (2011).

43. Evans, R. C. et al. The effects of NMDA subunit composition on calcium influx and spike timing-dependent plasticity in striatal medium spiny neurons. PLoS Comput Biol 8, e1002493 (2012).

44. Gerstner, W., Lehmann, M., Liakoni, V., Corneil, D. & Brea, J. Eligibility Traces and Plasticity on Behavioral Time Scales: Experimental Support of NeoHebbian Three-Factor Learning Rules. Front Neural Circuits 12, 53 (2018).

45. Magee, J. C. & Grienberger, C. Synaptic Plasticity Forms and Functions. Annu Rev Neurosci 43, 95–117 (2020).

46. Kwon, T., Sakamoto, M., Peterka, D. S. & Yuste, R. Attenuation of Synaptic Potentials in Dendritic Spines. Cell Rep 20, 1100–1110 (2017).

47. Cornejo, V. H., Ofer, N. & Yuste, R. Voltage compartmentalization in dendritic spines in vivo. Science 375, 82–86 (2022).

48. Bartol, T. M. et al. Nanoconnectomic upper bound on the variability of synaptic plasticity. Elife 4, e10778 (2015).

49. Motta, A. et al. Dense connectomic reconstruction in layer 4 of the somatosensory cortex. Science 366, (2019).

50. Reiner, A. Corticostriatal projection neurons – dichotomous types and dichotomous functions. Frontiers in Neuroanatomy vol. 4 Preprint at 10.3389/fnana.2010.00142 (2010).

51. Deng, Y. et al. Differential organization of cortical inputs to striatal projection neurons of the matrix compartment in rats. Front. Syst. Neurosci. 9, 51 (2015).

52. Cragg, B. Preservation of extracellular space during fixation of the brain for electron microscopy. Tissue and Cell vol. 12 63–72 Preprint at 10.1016/0040-8166(80)90052-x (1980).

53. Scheffer, L. K., Karsh, B. & Vitaladevun, S. Automated Alignment of Imperfect EM Images for Neural Reconstruction. arXiv [q-bio.QM] (2013).

54. Sato, M., Bitter, I., Bender, M. A., Kaufman, A. E. & Nakajima, M. TEASAR: tree-structure extraction algorithm for accurate and robust skeletons. in Proceedings the Eighth Pacific Conference on Computer Graphics and Applications 281–449 (IEEE Comput. Soc, 2000).

55. Mosteller, F. & Tukey, J. W. Data analysis, including statistics. Handbook of social psychology 2, 80–203 (1968).

56. Bellingham, M. C., Lim, R. & Walmsley, B. Developmental changes in EPSC quantal size and quantal content at a central glutamatergic synapse in rat. The Journal of Physiology 511, 861–869 (1998).

57. Bartos, M. et al. Fast synaptic inhibition promotes synchronized gamma oscillations in hippocampal interneuron networks. Proc Natl Acad Sci U S A 99, 13222–13227 (2002).

58. Kojima, S., Kao, M. H. & Doupe, A. J. Task-related ‘cortical’ bursting depends critically on basal ganglia input and is linked to vocal plasticity. Proc Natl Acad Sci U S A 110, 4756–4761 (2013).

59. Tanaka, M., Singh Alvarado, J., Murugan, M. & Mooney, R. Focal expression of mutant huntingtin in the songbird basal ganglia disrupts cortico-basal ganglia networks and vocal sequences. Proc Natl Acad Sci U S A 113, E1720–7 (2016).

60. Tønnesen, J., Katona, G., Rózsa, B. & Nägerl, U. V. Spine neck plasticity regulates compartmentalization of synapses. Nat Neurosci 17, 678–685 (2014).

61. Wolf, J. A. et al. NMDA/AMPA ratio impacts state transitions and entrainment to oscillations in a computational model of the nucleus accumbens medium spiny projection neuron. J Neurosci 25, 9080–9095 (2005).

62. Nisenbaum, E. S., Wilson, C. J., Foehring, R. C. & Surmeier, D. J. Isolation and characterization of a persistent potassium current in neostriatal neurons. Journal of Neurophysiology (1996) doi:10.1152/jn.1996.76.2.1180.

63. Gertler, T. S., Chan, C. S. & Surmeier, D. J. Dichotomous anatomical properties of adult striatal medium spiny neurons. J Neurosci 28, 10814–10824 (2008).

64. Al-Muhtasib, N., Forcelli, P. A. & Vicini, S. Differential electrophysiological properties of D1 and D2 spiny projection neurons in the mouse nucleus accumbens core. Physiol Rep 6, e13784 (2018).

65. Planert, H., Berger, T. K. & Silberberg, G. Membrane properties of striatal direct and indirect pathway neurons in mouse and rat slices and their modulation by dopamine. PLoS One 8, e57054 (2013).

66. Farries, M. A. & Perkel, D. J. Electrophysiological properties of avian basal ganglia neurons recorded in vitro. J Neurophysiol 84, 2502–2513 (2000).

67. Nisenbaum, E. S. & Wilson, C. J. Potassium currents responsible for inward and outward rectification in rat neostriatal spiny projection neurons. J Neurosci 15, 4449–4463 (1995).

68. Jahr, C. E. & Stevens, C. F. A quantitative description of NMDA receptor-channel kinetic behavior. J. Neurosci. 10, 1830–1837 (1990).

