## Supplementary Information for "An anatomical substrate of credit assignment in reinforcement learning"

### Supplementary Material

#### Supplementary Notes and Figures

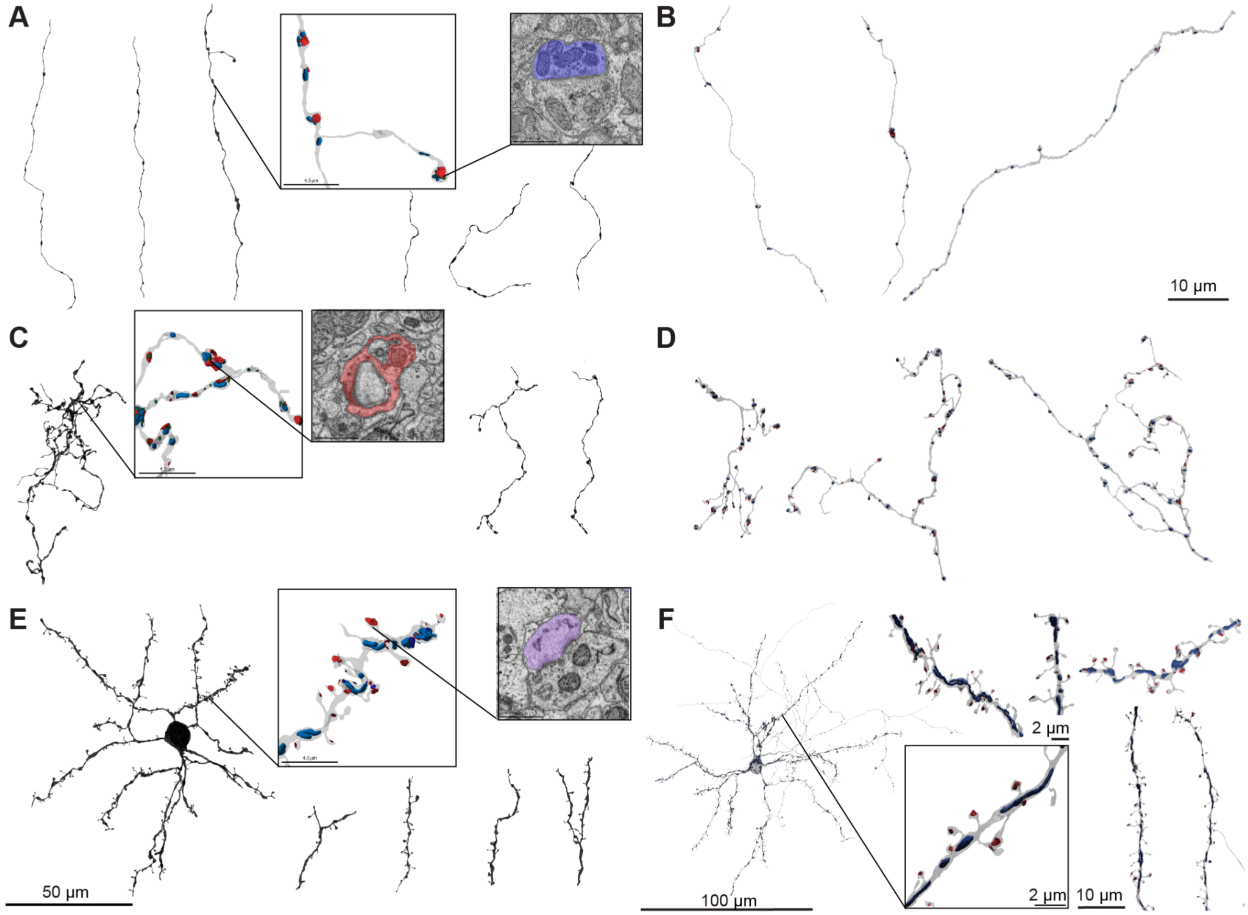


**Fig. S1 Examples of HVC, LMAN and MSN reconstructions**
(A) Example renderings of HVC axons in j0126. (B) Example renderings of HVC axons in j0251. (C) Example renderings of LMAN axons in j0126. (D) Example renderings of LMAN axons in j0251. (E) Example renderings of a MSN cell and MSN spiny dendrites in j0126. (F) Example renderings a MSN cell and MSN spiny dendrites in j0251. (A, C, E) are all on the same scale. (B, D) Examples are on the same scale. Minimum neurite length 50 µm skeleton path length. The zoomed insets show characteristics of the particular cell types, namely regular boutons for HVC axons, a perforated morphology of LMAN axons, and spiny dendrites of MSNs. Excitatory synapses (red), as determined by the synapse type classifier, mitochondria (blue), vesicle clouds (green). (B, D, F) show synapses in red and mitochondria in blue.

**Supplementary Note - Comparison of analyses with and without proofreading in j0126**

To assess the degree to which the proofreading of the automated reconstructions affected our conclusions, we created an automated connectome without manual healing of splits (reconnects) or false mergers. The resulting segmentation contained slightly more neurites (n=46,033 vs. 41,324), due to a higher split rate. This was not offset by the additional false mergers, and had a marginal effect on the overall neurite length distributions. We recreated key analyses regarding the model predictions, and found virtually no differences (Fig. S2). While our results demonstrate the dramatic progress of automated reconstruction methods for vEM data sets, we stress that manual proofreading still improved the quality of the data set.

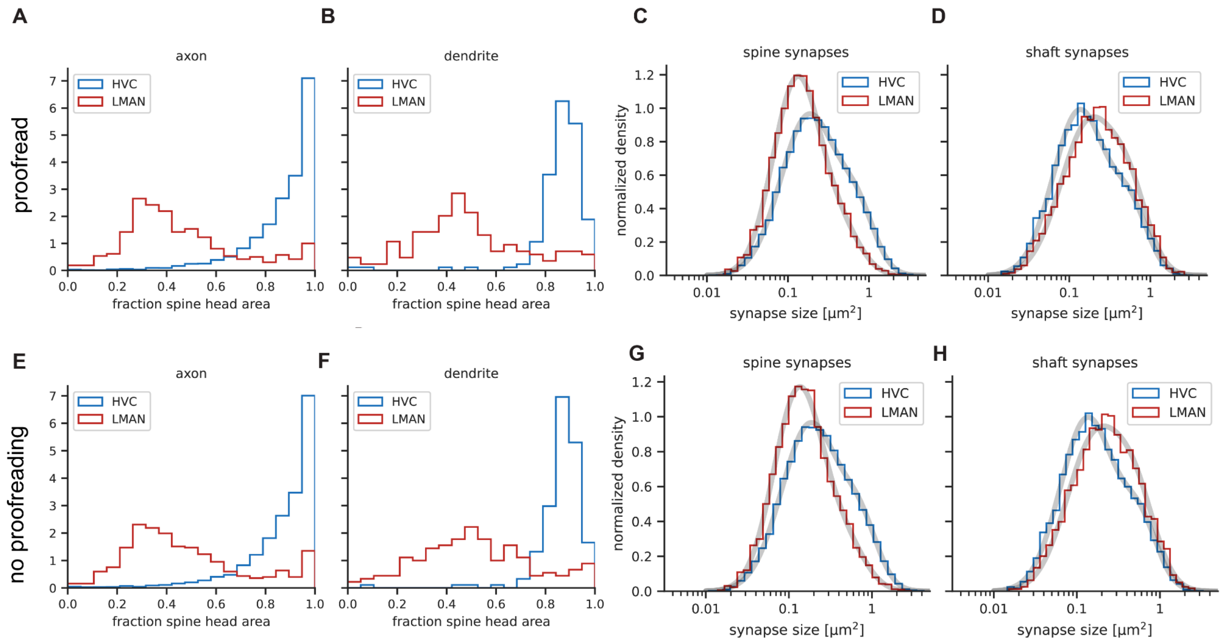


**Fig. S2: Spine / shaft preference in the j0126 dataset with and without manually corrected neuron reconstruction errors.**

(A-D) Analysis of spine / shaft preference of LMAN and HVC axons, as in Fig. 2 C-F, for the proofread version of j0126. (E–H). Same plots as in (A–D) from the j0126 data set, but without proofreading.

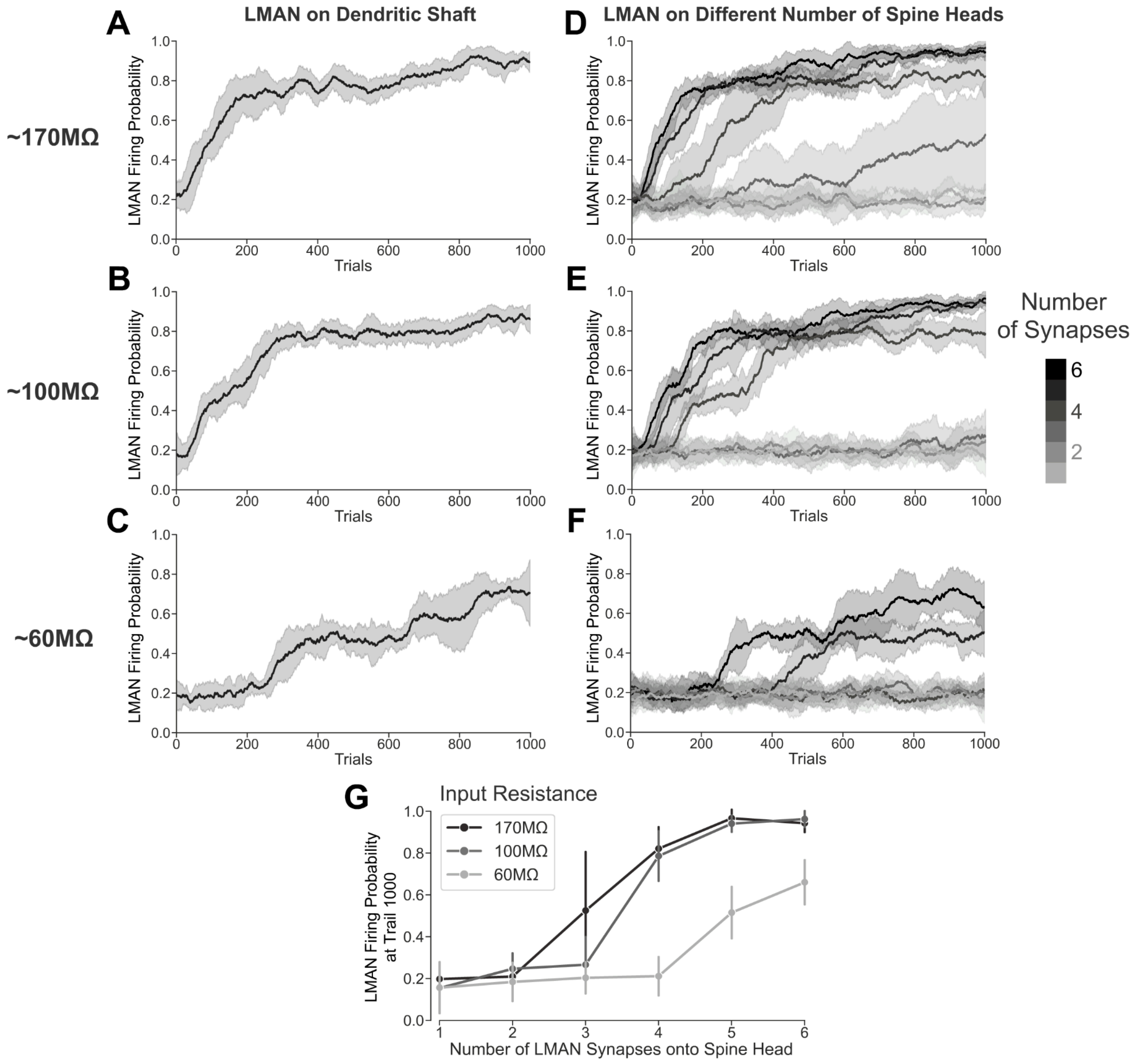


**Fig. S3 Performance throughout learning when the LMAN neuron synapses onto the dendritic shaft versus different numbers of spine heads, under different input resistances.**

(A-C) Performance throughout learning when the LMAN neuron synapses onto the dendritic shaft. (D-F) Performance throughout learning when the LMAN neuron synapses onto different numbers of spine heads. Each row represents one level of input resistance measured at the soma at -85 mV. (G) Performance at trial 1000 when the LMAN neuron synapses onto different numbers of spine heads, shown for 3 different input resistances. The shaded areas and vertical bars indicate the standard deviation across 10 repetitions.

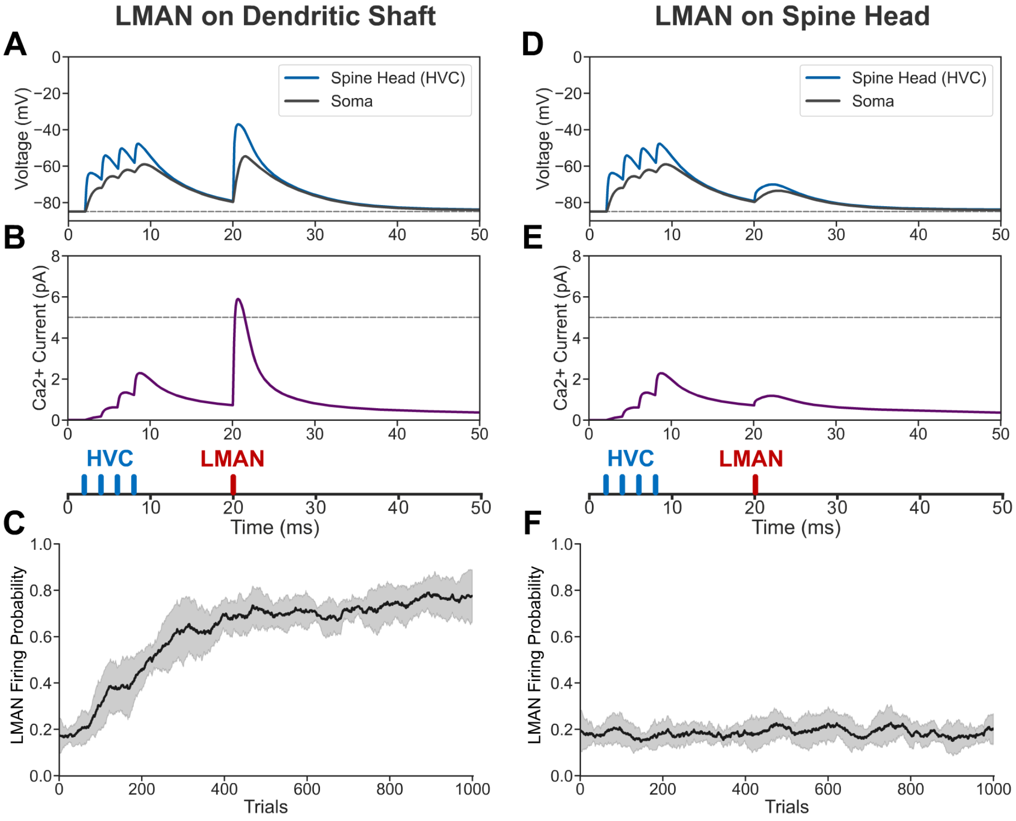


**Fig. S4 Results obtained when the MSN is modeled with synapses with the actual spatial distribution and actual morphology.**

(A-C) Results obtained if the LMAN neuron synapses onto the dendritic shaft. (D-F) Results obtained if the LMAN neuron synapses onto the spine head. The shaded areas indicate the standard deviation across 10 repetitions. As in the model of the main text that had equally spaced synapses of identical morphology, the model with synapse locations and morphologies taken from the actual reconstructed neuron again shows that there is learning only when the LMAN neuron synapses onto the dendritic shaft but not when it synapses onto a spine head.
